# BET bromodomain inhibitor HMBA synergizes with MEK inhibition in treatment of malignant glioma

**DOI:** 10.1101/2020.01.01.891739

**Authors:** Elisa Funck-Brentano, Dzeneta Vizlin-Hodzic, Jonas A. Nilsson, Lisa M. Nilsson

## Abstract

**Background:** BET bromodomain proteins regulate transcription by binding acetylated histones and attracting key factors for e.g. transcriptional elongation. BET inhibitors have been developed to block pathogenic processes such as cancer and inflammation. Despite having potent biological activities, BET inhibitors have still not made a breakthrough in clinical use for treating cancer. Multiple resistance mechanisms have been proposed but thus far no attempts to block this in glioma has been made.

**Methods:** Here, we have conducted a pharmacological synergy screen in glioma cells to search for possible combination treatments augmenting the apoptotic response to BET inhibitors. We first used HMBA, a compound that was developed as a differentiation therapy four decades ago but more recently was shown to primarily inhibit BET bromodomain proteins. Data was also generated using other BET inhibitors.

**Results:** In the synergy screen, we discovered that several MEK inhibitors can enhance apoptosis in response to HMBA in rat and human glioma cells in vitro as well as in vivo xenografts. The combination is not unique to HMBA but also other BET inhibitors such as JQ1 and I-BET-762 can synergize with MEK inhibitors.

**Conclusions:** Our findings validate a combination therapy previously demonstrated to exhibit anti-cancer activities in multiple other tumor types but which appears to have been lost in translation to the clinic.

## 1. Introduction

Before the discovery of oncogenes the concept of cancer cell differentiation therapy was explored therapeutically, in part based on early observations that dimethylsulfoxide (DMSO) can cause differentiation of Friend virus induced mouse erythroleukemia (MEL) cells into hemoglobin producing red blood cells (1). Efforts to produce more potent cancer differentiation compounds generated two molecules that were tested in the clinic, hexamethylene bisacetamide (HMBA) and suberoylanilide hydroxamic acid (SAHA, later renamed to vorinostat) (2, 3). Whereas SAHA was found to inhibit histone deacetylases (HDACs) 1-3 and made it to clinical approval for cutaneous T-cell leukemia, HMBA neither inhibits HDACs nor received clinical approval, and its target was unknown for forty years (4, 5). Recently, however, we discovered that HMBA is a bromodomain and extra-terminal domain (BET) inhibitor, with highest binding affinity for bromodomain 2 (BD2) of BET proteins BRD2, BRD3 and BRD4 while also inhibiting the bromodomain of histone acetyltransferase P300 (6). The structure of HMBA largely resembles that of an acetylated lysine, explaining the mode of action.

Although HMBA was likely the first anti-cancer compound used in the clinic that inhibited BET bromodomain proteins, the concept of BET inhibitors (BETis) were largely popularized with the development of the low nanomolar BETis JQ1 and iBET-151 (7, 8). The mechanism of action of these compounds involves inhibition of transcriptional elongation (9, 10). Albeit that MYC transcription is frequently suppressed, all effects of BETis are not dependent on MYC suppression (11, 12). Most of the clinical studies using HMBA and other BETis have focused on hematological malignancies and less is known about the effect of this class of compound in solid tumors such as glioma. Hematological malignancies respond to BETis in vitro by cell cycle arrest, differentiation and apoptosis, whereas glioma cells undergo cell cycle arrest and differentiation and to a lesser extent apoptosis (13–15). Importantly for glioma treatments, the clinical BET inhibitor OTX015 has been shown to pass the blood-brain-barrier (16).

If BETis are to work in the clinic against solid tumors including glioma, then the predominant effect of BETis should be apoptosis. So far, BETis have not convincingly shown this effect as single agents in solid tumors. Here we use the C6 rat glioma model system to study means to activate cell death in BETi-treated cells. We demonstrate that the MAPK pathway is critical for maintaining viability of HMBA-treated C6 cells and demonstrate synergy between HMBA and MEK inhibitors in vitro and in mouse xenograft experiments using C6 cells and human primary glioma sphere cultures.

## 2. Results

To study the effect of HMBA in glioma we used the rat glioma cell line C6. Treatment of these cells for 72 hours blocked cell proliferation (Figure 1A) but did not induce apoptosis, as assessed by flow cytometry of sub-G1 DNA content (Figure 1B-C). We therefore conclude that C6 glioma cells primarily respond to HMBA by growth inhibition. Since apoptosis is the preferred mode of effect of cancer treatment we hypothesized that a signaling pathway targeted by drugs could be used by the cell to maintain viability upon HMBA treatment. We therefore screened a library of 226 compounds (Supplemental Table S1) either approved for clinical use or under various stages of clinical development. Comparing the effect of monotherapy of HMBA, with monotherapy of either library compound alone or with combination therapy of HMBA and library compound, we identified compounds that displayed synergistic effects together with HMBA, of which three were MEK inhibitors (Figure 1D).

**Figure 1.**
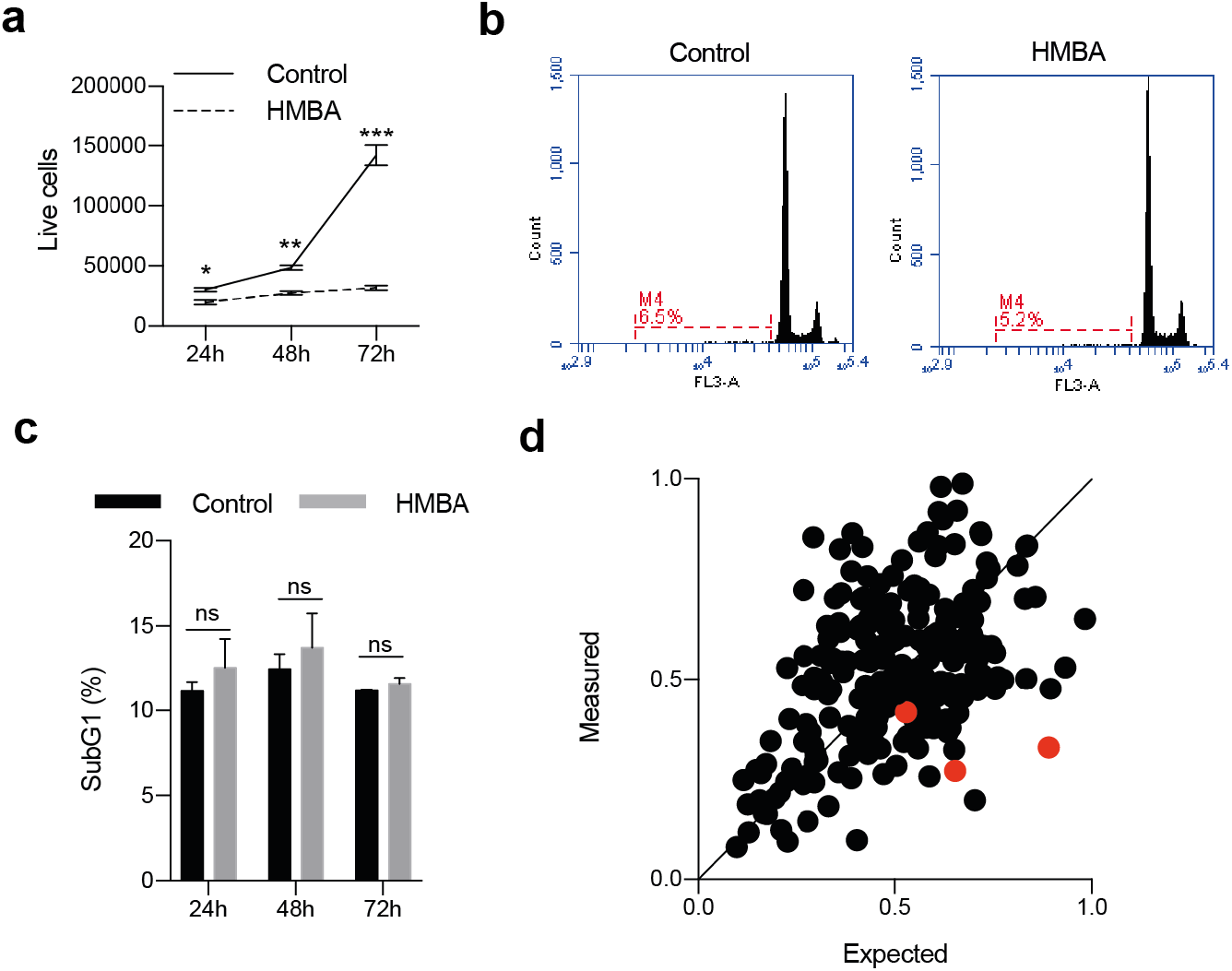
HMBA evokes primarily growth arrest in C6 glioma cells. a) Cell counts using trypan blue and b) DNA histograms of 7-AAD-stained nuclei quantifying the sub-G1 content together indicate that the primary response to HMBA-treatment in C6 glioma cells is growth arrest. c) Quantification of cells with less than diploid DNA content in b). d) Plot summarizing the results from the pharmacological screen of HMBA in combination with 226 different compounds. The three red dots indicate the three MEK inhibitors which all fall below the line of equal measured and expected.

Currently, two MEK inhibitors are FDA approved for use in melanoma but none are used for treatment of glioma. We repeated the results from the library screen using the FDA-approved MEK-inhibitor trametinib (GSK1120212) in a clonogenic assay (Figure 2A). The lack of long-term growth and induction of cell death was revealed by an ATP/luciferase-based viability assay (Figure 2B) and flow cytometric analysis of sub-G1 DNA content (Figure 2C). This cell death was likely mediated by caspases since the sub-G1 content of the cells could be rescued by the pan-caspase inhibitor Q-VD-OPH. Furthermore, the synergistic effects of dual BET and MEK inhibition could be reproduced using other MEK inhibitors (TAK-733 and AZD8330, but not binimetinib) and BETi (JQ1 or iBET-762; Figure 2D-E and Supplemental Figure S1A-B).

**Figure 2.**
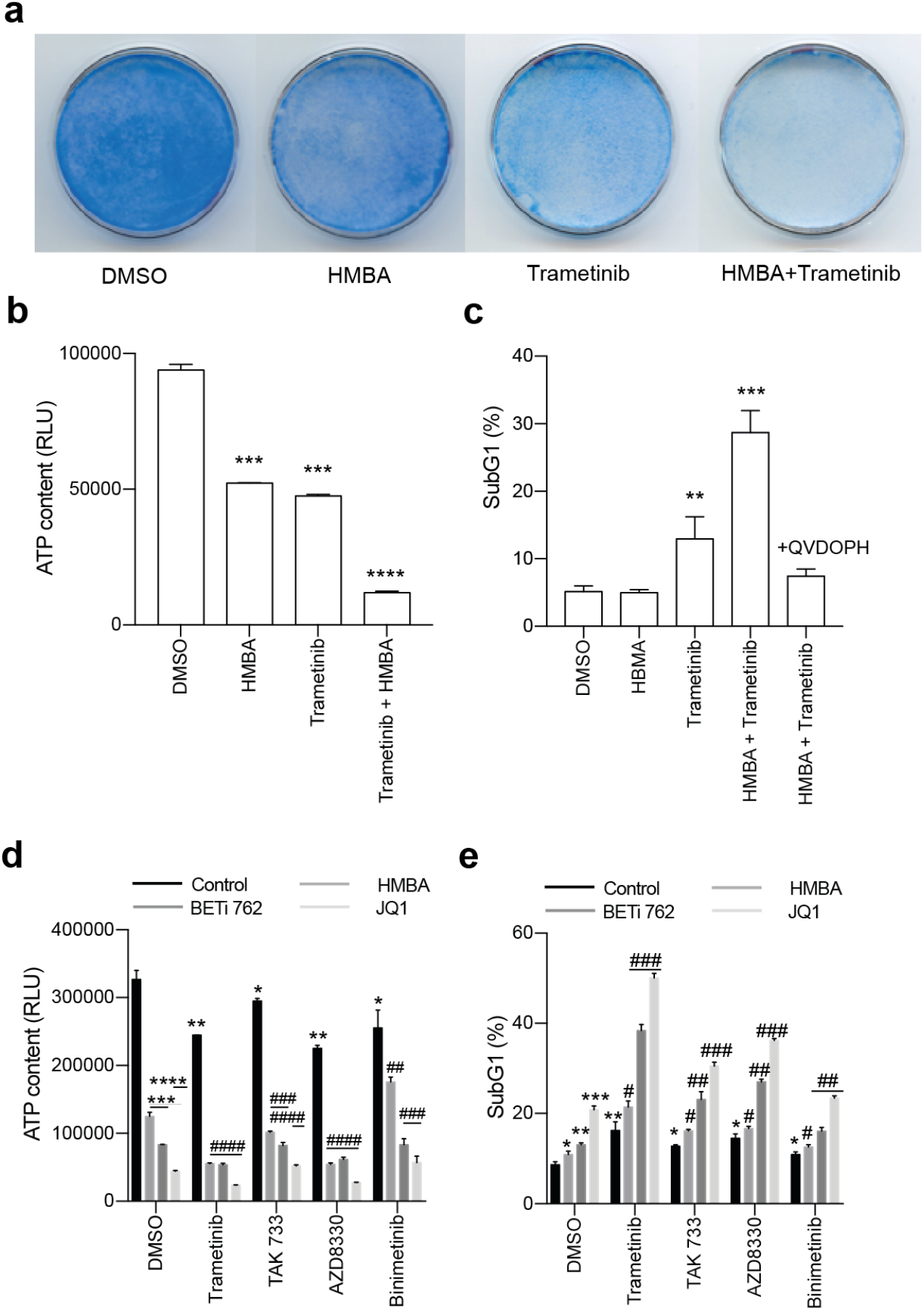
Combination of BET inhibitors and MEK inhibitors enhance cell death in C6 rat glioma cells. a) Clonogenic assay of C6 treated with HMBA, trametinib or the combination of both. b) Cell viability of single treatment or combination using Cell Titer Glo. The dotted indicates the expected value of an additive effect of the combination c) Percent of cells with less than diploid DNA content (sub-G1). The high rates in combination treatment could be suppressed with pan-caspase inhibitor Q-VD-OPH, suggesting apoptosis. d) and e) Viability and sub-G1 assessments of combinations of BET inhibitors HMBA, JQ1 and I-BET762 together with MEK inhibitors trametinib, TAK733, AZD8330 and binimetinib. Singe asterisks or hash signs indicate significant values of p<0.05, double signs are p<0.01, triple signs are p<0.001 and quadruple signs are p<0.0001.

Earlier studies had indicated that HMBA could more efficiently induce differentiation of a vincristine-resistant leukemia cell line (17). This suggested that HMBA possibly could interfere with drug resistance pump such as p-glycoprotein (ABCB1 or MDR1) but such a link could not be established. On the other hand, trametinib had previously been shown to be a substrate of p-glycoprotein (p-gp) (18) so we reasoned that p-gp could be involved in the synergy in C6 glioma cells. Indeed, as previously shown (19), C6 cells were highly effective in pumping out the substrate rhodamine 123, and this activity was blocked by the ABCB1/ABCG2 inhibitor elacridar (Figure 3A). Interestingly, also trametinib – but not the other MEK inhibitors tested – could inhibit pumping of rhodamine 123 but only occasionally at 1 μM and more prominently at > 1μM (Supplemental Figure 3A-B). Blocking of pumps with elacridar reduced the concentration needed to inhibit ERK phosphorylation in C6 cells (Figure 3B). However, the fact that elacridar neither synergized with trametinib nor with HMBA or JQ1 (Figure 3C-D) made it unlikely that BETis synergize with MEK inhibitors because of regulation of p-gp or other drug pumps. Rather, as HMBA and JQ1 synergize with other MEK inhibitors (Figure 2D-F) – which were not p-gp inhibitors (Supplemental Figure 3A-B) – and also with the ERK inhibitor SCH772984 (Figure 3E-G), suggests that the MAPK pathway maintains viability of BETi-treated C6 glioma cells.

**Figure 3.**
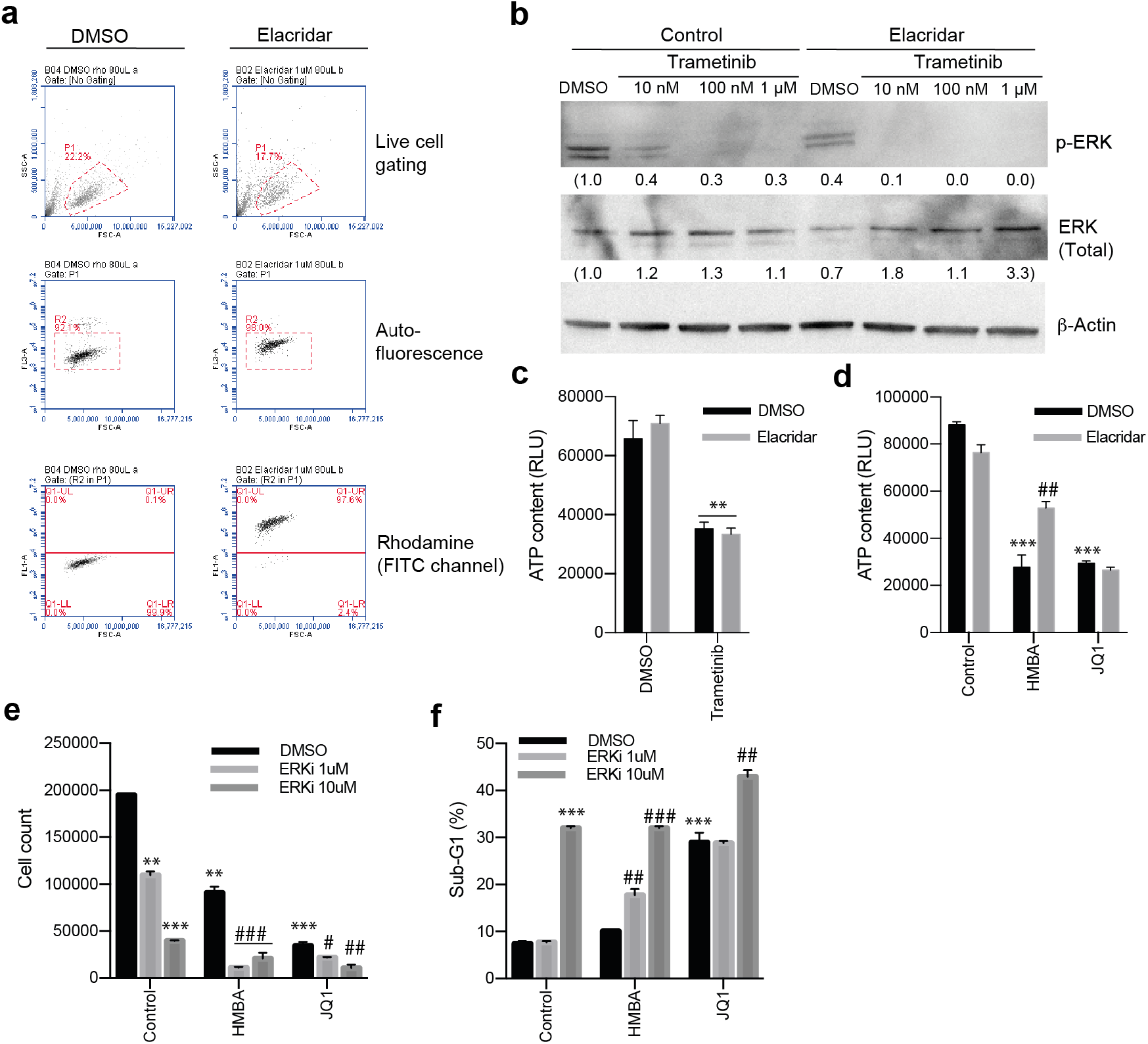
Inhibiting p-gp activity with elacridar affects trametinib activity but does not synergize with trametinib to kill C6 cells. a) Flow cytometry analysis for p-gp activity showing p-gp substrate Rhodamine 123 being pumped out of cells (DMSO, bottom panel whereas inhibiting p-gp with elacridar blocks this pumping (Elacridar, bottom panel). b) Blocking p-gp with elacridar enhanced the activity of trametinib as judged by lowered P-ERK on Western blot. Values of relative expression to actin and the control, assessed by densitometry, is below images. Uncropped images are in Supplemental Figure S2. c) Viability assay showing that inhibiting p-gp with elacridar does not enhance killing by trametinib. d) Viability assay of combination treatments with elacridar and HMBA or JQ1. e) and f) Viability and sub-G1 assessments of combinations of BET inhibitors HMBA and JQ1 together with ERK inhibitor SCH772984.

Next, we investigated if HMBA and trametinib could block tumor growth in vivo. Immunocompromised NOG mice were transplanted with C6 cells subcutaneously, and when tumors were palpable they were randomized to treatment either with normal food or food containing trametinib and/or normal drinking water or drinking water supplemented with HMBA. Tumors in mice treated with HMBA in drinking water or trametinib in the food grew significantly slower than tumors growing in control mice and in HMBA/trametinib-treated mice tumor growth was robustly suppressed resulting in four-fold longer survival (Figure 4A-B).

**Figure 4.**
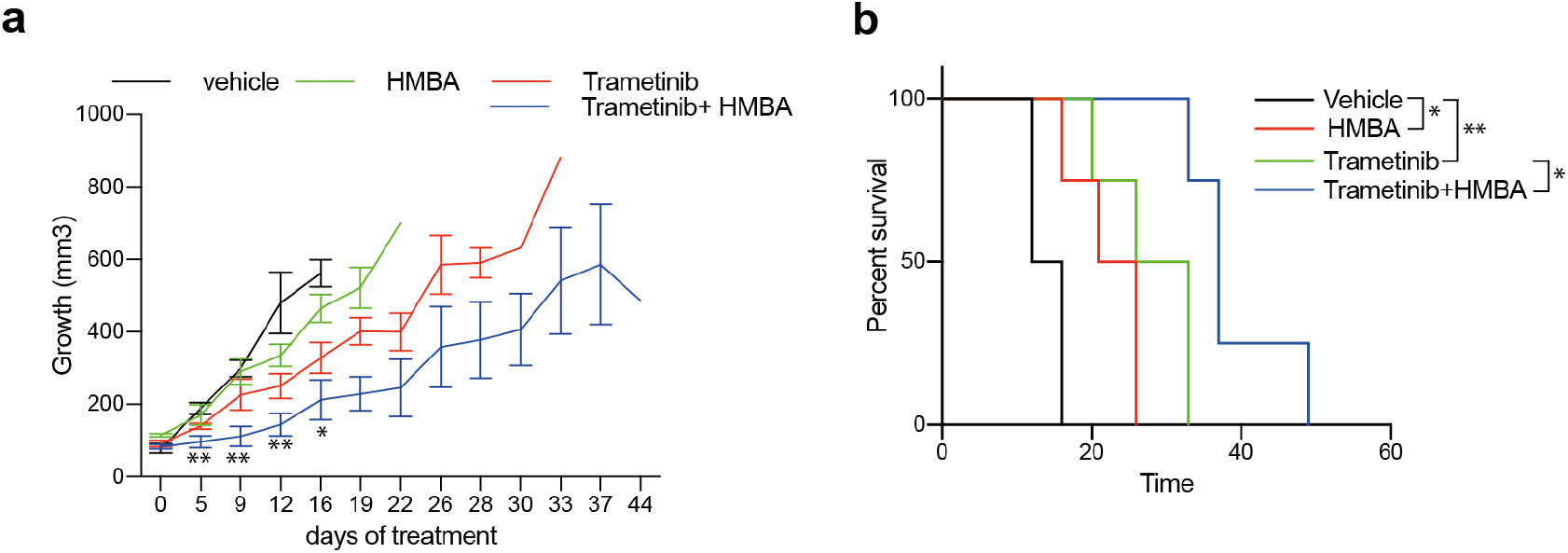
Treatments in vivo of C6 glioma with HMBA (2.5 % HMBA in drinking water), trametinib (0.5 mg/kg mouse mixed in food) or combination. a) Tumor volumes over time with respective treatment. Significance of curve comparisons (asterisks) are made for vehicle and trametinib+HMBA treated mice. b) Survival curve indicating the elapsed time for tumors in the different treatment to reach ethical size limit, n=4 in each group.

To gain insight into whether or not the described synergy effect of BET and MEK inhibition would also impact on human glioma we treated four patient-derived glioma sphere cultures with HMBA and trametinib. Three out of the four cell lines had some response to trametinib in short-term sphere culture but only one out of the four cell lines, NCH421K, was sensitive to the combination, suggesting that multiple pathways maintain viability of human glioma cells treated with HMBA (Figure 5A). However, long-term adherent culture of NCH644 and NCH690 revealed sensitivity to the combination (Figure 5B). Nevertheless, treatment of mice bearing NCH421K tumors with HMBA water and trametinib food suppressed growth (Figure 5C). Trametinib has been associated with induction of kinase activities in triple-negative breast cancer cells through enhancer remodeling (20). Presumably, this could help the cells survive MEK inhibition, which would be perturbed by BETi treatment if these kinases rely on BET protein-regulated processes for expression. In order to investigate if this also holds true in glioma, we performed phosphokinase arrays on two of the human glioma lines, NCH644 and NCH690. The analysis included 43 phosphorylation sites of known kinases. After 24 hours’ treatment, there were minor effects on kinase activities in the two cell lines (Figure S3). The graphs display the ratios of trametinib vs vehicle of each cell line. Reassuringly, ERK phosphorylation and phosphorylation of the ERK target CREB was inhibited in both cell lines, confirming the activity of the MEK inhibitor. Glioma line NCH690 exhibited a general downregulation of all kinase activities tested in the assay, in accordance with the overt sensitivity of this cell line to monotherapy with trametinib (Figure 5B). The NCH644 line, on the other hand, was less affected by monotherapy with trametinib (Figure 5B) and phosphorylation of for example c-Jun, FYN and PRAS40 was induced by trametinib (Figure S4). Collectively, our data does not provide a consistent view on changes of the phospho-proteome in glioma cells treated with trametinib, besides inhibition of ERK phosphorylation.

**Figure 5.**
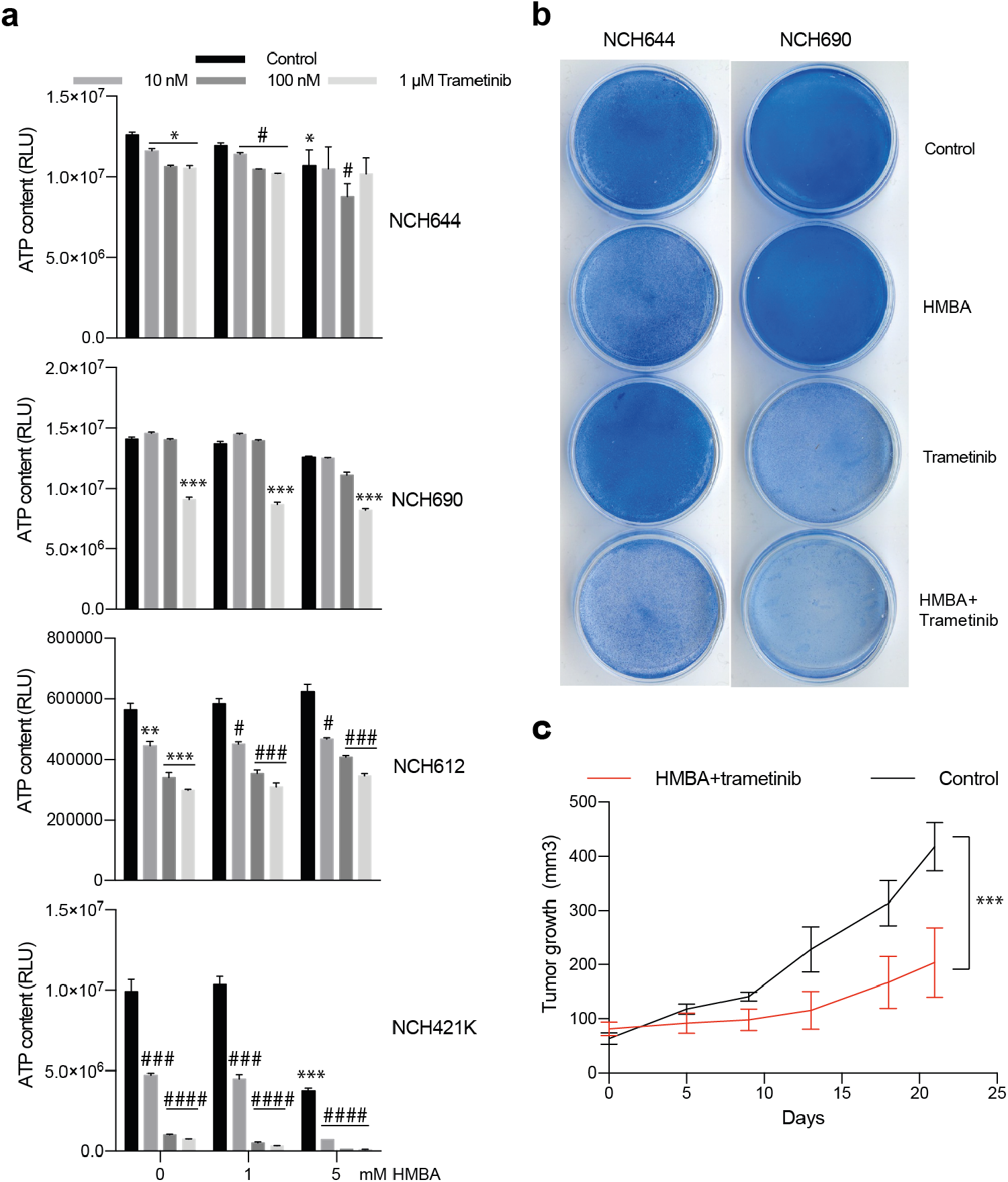
Human glioma cells are sensitive to combination of HMBA and trametinib. a) Viability assay of four human glioma cell lines treated for three days as spheres with HMBA and varying concentrations of trametinib. b) Clonogenic assay of cell lines NCH644 and NCH690 growing adherently on plastic showing potency of combination treatment. c) Tumor growth in vivo of NCH421K cells growing subcutaneously on NOG mice and treated with combination of HMBA+trametinib.

## 3. Discussion

In the present study we have identified means to enhance efficacy of BETis in models of glioma. Several BETis have already entered clinical trials, e.g. HMBA, OTX015 and ABBV-075 (21–23), but thus far the therapeutic effect of these inhibitors as monotherapies have been sparse. Our findings that MEK inhibitors, which are already available in clinical use, could synergize with BETis is therefore of clinical interest. Notably though, the synergistic effect of simultaneous targeting with BET and MEK inhibitors has also been observed in a broad set of tumor types (20, 24–28). The sensitivity has been correlated to certain mutational states, like Suz12 loss in malignant peripheral nerve sheath tumors (24) which leads to an epigenetic switch from histone methylation to histone acetylation, rendering the tumors sensitive to BET inhibitor JQ1. Another study demonstrated that resistance to MEK inhibitors associated with BRD4-induced enhancer formation, which could be inhibited by JQ1 (20). Although the combination therapy can show effects in many tumor types it is not certain that the mechanism will be identical in all affected tumor types since the transcriptional effects of BET inhibition is very pronounced.

MEK/BET combination inhibition can suppress MAPK and checkpoint inhibitor resistant melanoma in animal models including those exhibiting NRAS mutations (27). Although BET and MEK inhibitors would be expected to have effects on normal lymphocytes as well, the combination had activity also in immune competent mice and did not impair immune cells. However, in these experiments, checkpoint inhibitors were not given which could explain the general insensitivity of the non-dividing immune cells to BET/MEK combination treatment.

We have previously published data demonstrating that BETis act as what historically was referred to cancer differentiating agents (6, 12). Tumor cell differentiation therapies held great promise during the 1980’s and 1990’s, but did not render any clinically approved therapies for solid tumors. The vast literature, including our study, on combining BET inhibitors with MAPK inhibitors, could be a solution to enhancing the effects of previously tested differentiation therapies. Glioma patients have very few viable treatment options for advanced disease and therefore could participate in phase 1 studies on the combination between BETis and e.g. trametinib. A possible challenge may be that trametinib appears to be a substrate of drug pumps (18) but this has to be validated in the clinic.

## 4. Materials and Methods

### 4.1. Chemicals

HMBA and Rhodamine 123 was purchased from Sigma-Aldrich. A collection of 226 anti-cancer compounds under clinical development or in clinical use as well as AZD8330, I-BET-762, trametinib, TAK733, binimetinib, elacridar and ERK inhibitor SCH772984 were all from Selleck Biochemicals. The (+)-enantiomer of JQ1 was purchased from Cayman chemicals.

### 4.2. Cell culture

The rat glioma cell line C6 was bought from Cell Line Service (CLS) and grown in RPMI-1640 supplemented with 10% FBS, GlutaMAX and antibiotics. The human glioma sphere cultures NCH412K, NCH612, NCH644 and NCH690 were form CLS and were cultured according to the company’s recommendations in glioma sphere medium MG43 (CLS) as spheres or adherent cultures using laminin-coated plastic dishes. Viability after treatments was analysed with Cell Titer Glo (Promega), or Coomassie-staining of cells grown for clonogenic assay.

### 4.3. Mouse experiments

All animal experiments were performed in accordance with regional/local animal ethics committee approval (approval number 36/2014). C6 or NCH412K cells were injected subcutaneously onto the flanks of immunocompromised, non-obese severe combined immune-deficient interleukin-2 chain receptor γ knockout mice (NOG mice; Taconic, Denmark). Tumors were measured with caliper at regular time points and tumor volumes were calculated using the formula: tumor volume (mm3)=(length(mm)) × (width(mm))2/2. Treatments were started when the tumors were actively growing, judged by increasing volumes on repeated caliper measurements. Trametinib was mixed in the chow at 2.5 mg/kg giving an approximate dose of 0.5mg/kg mouse per day. HMBA was given in drinking water as 2.5% HMBA, 0.33g/L bicarbonate, 2% sucrose. Vehicle was given as 0.33 g/L bicarbonate, 2% sucrose. Mice were sacrificed and tumors were harvested before or when tumors reached ethical size limit.

### 4.4. Cell cycle analysis

One million cells per mL were lysed and stained for 30 minutes at 37°C in modified Vindelöv’s solution (20 mM Tris, 100 mM NaCl, 1 μg/mL 7-AAD, 20 μg/mL RNase, and 0.1 % NP40 adjusted to pH 8.0) followed by analysis of DNA content using the FL3 channel (linear mode and cell cycle) or FL3 channel (logarithmic mode and apoptosis) with a BD Accuri C6 flow cytometer.

### 4.5. Western blot

For western blot analysis of protein expression, cell pellets or tumor pieces were lysed in lysis buffer (50 mM HEPES pH 7.5, 150 mM NaCl, 1 mM EDTA, 2.5 mM EGTA, 0.1 % Tween-20, 1 x HALT protease and phosphatase inhibitors (Thermo Scientific)) on ice. After sonication and clearing of lysates, protein was determined using Bio-Rad Protein Assay Dye reagent (Bio-Rad). A total of 50 μg of protein was resolved on 4–20% Mini-PROTEAN TGX gels (Bio-Rad) and transferred to nitrocellulose membrane (Protran, GE Healthcare Bio-Sciences). Membranes were stained with Ponceau S red dye to verify equal loading. All subsequent steps were carried out in TBS-Tween (10 mM Tris-HCl, pH 7.6, 150 mM NaCl, and 0.05 % Tween-20) containing 5% bovine serum albumin for antibody incubations. Antibodies against total ERK and phosphorylated ERK were from Cell Signaling, beta-Actin was from Sigma. For phosphorylation site detection the Proteome profiler human phospho-kinase array kit (R&D Systems) was used according to manufacturer’s instructions. Lysates were prepared the same way as described above and 200 μg total protein was incubated with each membrane set. The signals were quantified using densitometry.

### 4.6. Analysis of pump activity

C6 cells were treated for 48 hours with indicated inhibitor, after which they were further treated in the presence of 200 ng/mL Rhodamine 123 for 60 min. After incubation, the cells were washed with PBS and cultured for another 90 min in fresh medium with continued treatment but in the absence of Rhodamine 123. Elacridar was added (1uM) to block pumping of Rhodamine 123. Cells were harvested and resuspended in PBS and analyzed with a BD Accuri C6 flow cytometer.

### 4.7. Statistical analysis

Graphs were generated using GraphPad Prism, error bars on tumor growth curves are shown as standard error of mean (SEM), and error bars on cell experiments are shown as standard deviation (SD). Statistical significance was assessed by Student’s T test and significant values compared to vehicle are indicated by asterisks whereas significant values compared to relevant monotherapy in combination experiments are indicated by hash signs. Singe asterisks or hash signs are p<0.05, double signs are p<0.01, triple signs are p<0.001 and quadruple signs are p<0.0001. Survival curve analysis for in vivo experiments was performed using the log-rank (Mantel-Cox) test in Graph Pad Prism (GraphPad Software).

## 5. Conclusions

The present study confirms in an additional cancer type that targeting BET bromodomain protein and MEK is more effective than monotherapies of both inhibitors. We propose the initiation of a basket clinical trial for patients with solid tumors that have failed targeted therapies and/or immunotherapies.

## Supplementary Materials

The following are available: Figure S1-4 and Table S1-2.

## Author Contributions

The following were made: Conceptualization, J.A.N. and L.M.N.; methodology, E.F.B, and L.M.N.; data curation, E.F.B., J.A.N., L.M.N.; writing—original draft preparation, E.F.B., J.A.N., L.M.N.; writing—review and editing, E.F.B, D.V.H., J.A.N., L.M.N; supervision, J.A.N. and L.M.N.

## Acknowledgments

We thank Sofia Stenqvist and Carina Karlsson for technical support.

