## Supplementary material for "BET bromodomain inhibitor HMBA synergizes with MEK inhibition in treatment of malignant glioma": Figure S

### Supplemental Figures

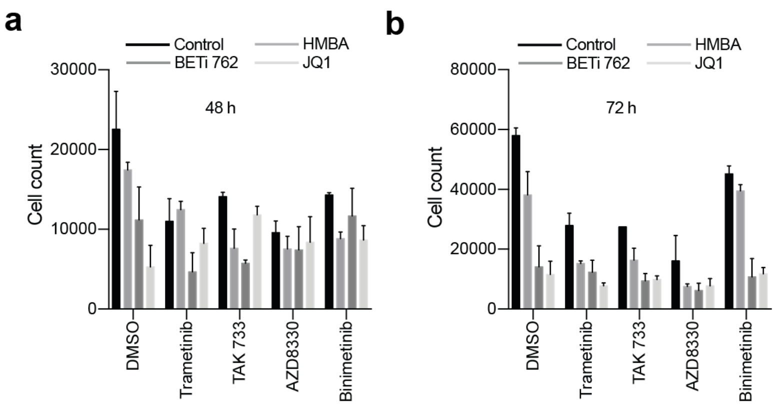


**Figure S1.** Combination treatment with different BET and MEK inhibitors synergize in killing of C6 cells. a) Cell counts after 48 hours treatment with four different MEK inhibitors in combination with three different BET inhibitors. b) Cell counts after 72 hours with the same compounds as in a).

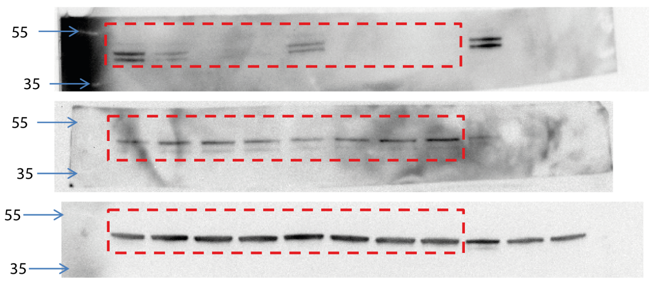


**Figure S2.** Related to figure 3: uncropped immunoblot images.

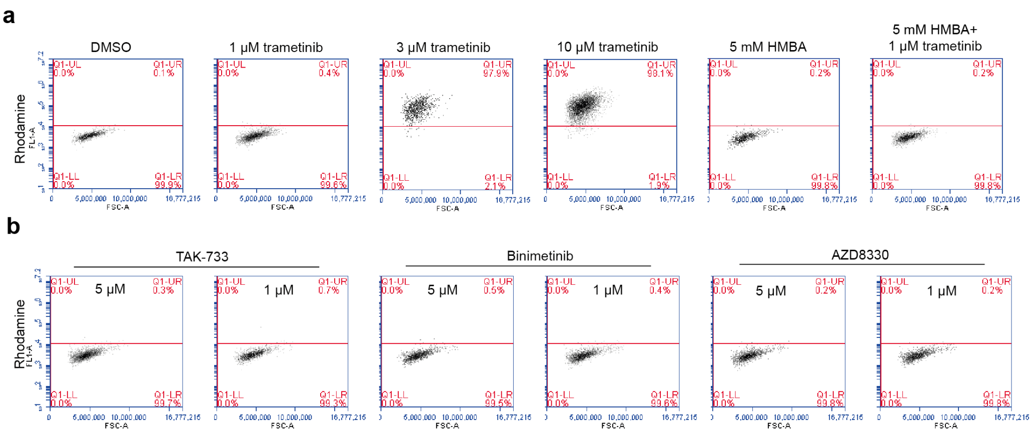


**Figure S3.** a) Trametinib can block p-gp activity of C6 cells judged by pumping of Rhodamine 123. a) Flow cytometry analysis of the p-gp substrate Rhodamine 123 over time in the presence of varying concentrations of trametinib or 5mM HMBA or combination of 5mM HMBA+1μM trametinib. b) Flow cytometry analysis of p-gp substrate Rhodamine 123 in the presence of MEK inhibitors TAK733, binimetinib and AZD8330.

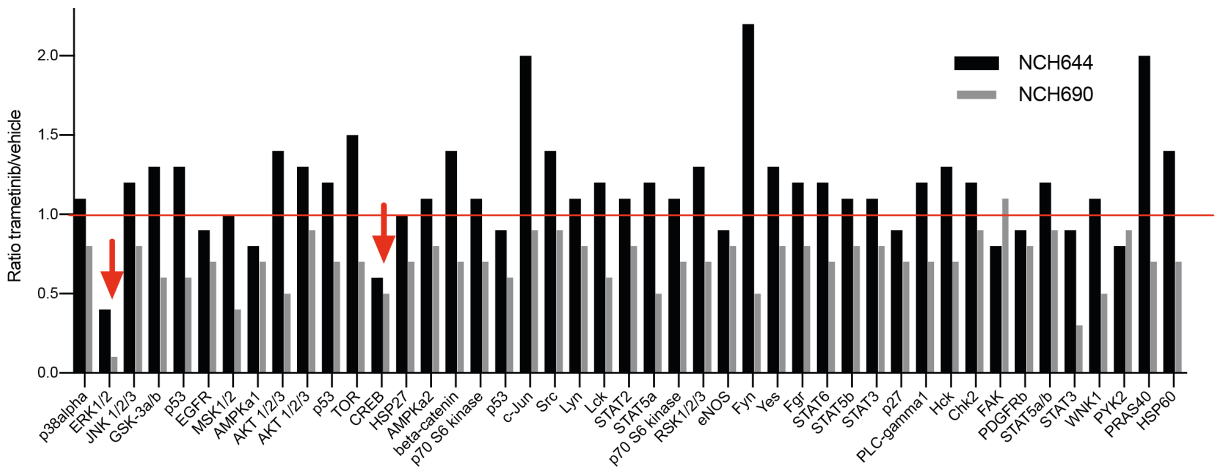


**Figure S4.** Effects on the phospho-proteome of trametinib in NCH644 and NCH690 cells. Phophorylation of indicated proteins were determined using a phosphoprotein membrane array. Phosphorylations were quantified using densitometry and values of trametinib-treated cells were divided by values of vehicle-treated cells (raw data are present in Supplemental Table S2). Arrows indicate known MEK/ERK target proteins.

**Supplemental Tables**

**Table S1.** All 226 compounds either alone or in combination with HMBA in C6 cells. Plates were stained with Coomassie and absorbances were read. Values were compared to vehicle control and HMBA alone.

**Table S2.** Densitometric measurements of phospo-protein array analysis of NCH644 and NCH690 cells treated with vehicle or 1 μM trametinib.
